# *APOE4* impacts cortical neurodevelopment and alters network formation in human brain organoids

**DOI:** 10.1101/2024.10.07.617044

**Authors:** Karina K. Meyer-Acosta, Eva Diaz-Guerra, Parul Varma, Adyasha Aruk, Sara Mirsadeghi, Aranis Muniz Perez, Yousef Rafati, Ali Hosseini, Vanesa Nieto-Estevez, Michele Giugliano, Christopher Navara, Jenny Hsieh

## Abstract

Apolipoprotein E4 (*APOE4*) is the leading genetic risk factor for Alzheimer’s disease. While most studies examine the role of *APOE4* in aging, imaging, and cognitive assessments reveal that *APOE4* influences brain structure and function as early as infancy. Here, we examined human-relevant cellular phenotypes across neurodevelopment using induced pluripotent stem cell (iPSC) derived cortical and ganglionic eminence organoids (COs and GEOs). In COs, we showed that *APOE4* decreased BRN2+ and SATB2+ cortical neurons, increased astrocytes and outer radial glia, and was associated with increased cell death and dysregulated GABA-related gene expression. In GEOs, *APOE4* accelerated maturation of neural progenitors and neurons. Multi-electrode array recordings in assembloids revealed that *APOE4* disrupted network formation and altered response to GABA, resulting in heightened excitability and synchronicity. Together, our data provides new insights into how *APOE4* may influence cortical neurodevelopmental processes and network formation in the human brain.

## Introduction

Alzheimer’s disease (AD) is the leading form of dementia causing progressive cognitive decline, with sporadic AD encompassing 95% of all cases (Harman, 2006). Apolipoprotein E4 (*APOE4*) is the leading genetic risk factor for sporadic AD, increasing risk 3-12-fold, with females demonstrating a greater risk (Karch et al., 2014; Liu et al., 2010; Verheijen and Sleegers, 2018). AD is characterized by amyloid β (Aβ) plaques and intracellular neurofibrillary tangles comprised of hyperphosphorylated tau protein (p-tau) (2023 Alzheimer’s disease facts and figures, 2023). These pathologies develop decades prior to cognitive decline, yet the early phenotypes preceding AD pathology are unknown (Holtzman et al., 2011).

Human *APOE4* carriers demonstrate altered brain structure and cognitive deficits as early as infancy. Imaging studies in infants have shown that *APOE4* reduces gray matter volume (GMV) and regional myelin water fractionation (MWF) in the precuneus, posterior/middle cingulate, lateral temporal, and medial occipitotemporal regions (preferentially affected by AD) while increasing GMV and MWF in the parietal lobe and frontal cortex (Dean et al., 2014; Knickmeyer et al., 2014). In children, *APOE4* decreases hippocampal volume and cortical thickness but increases frontal cortex volume. Cognitively, *APOE4* infants and toddlers score higher on early mental development tests, but *APOE4* children score lower on IQ, attention, and working memory tests (Chang et al., 2016; Remer et al., 2020; Reynolds et al., 2019; Wright et al., 2003). These studies suggest that *APOE4* influences cognition and brain structure well before the development of AD, potentially extending to neurodevelopment.

In *APOE4* mouse models, regional hyperexcitability, memory impairment, and altered adult neurogenesis are observed. Aged *APOE4* mice have increased excitatory neuron firing rate, synchronicity, and reduced responsiveness to inhibitory GABA inputs resulting in neuronal hyperexcitability (Area-Gomez et al., 2020). GABAergic signaling impairment or GABA neuron reduction is a common phenotype in *APOE4* mice. In fact, increasing GABAergic signaling chemically or by interneuron transplantation is sufficient to restore cognition, adult neurogenesis, and other AD-relevant phenotypes (Gillespie et al., 2016; Jang et al., 2023; Knoferle et al., 2014; Li et al., 2009; Montagne et al., 2021; Tong et al., 2016). This data suggests that *APOE4*-mediated loss of GABAergic inhibitory action affects cognition and adult neurogenesis.

Adult neurogenesis is decreased during early stages of AD in post-mortem brains, suggesting the observed influence of *APOE4* on adult neurogenesis in mice might be relevant to AD (Boldrini et al., 2018; Moreno-Jimenez et al., 2019; Tobin et al., 2019). While it is technically challenging to fully characterize the role of human adult neurogenesis, new neurons contribute to several cognitive processes in mice and are severely affected in AD (Lupo, 2023). ApoE has been shown to regulate both developmental and postnatal neurogenesis in mice, and both *APOE4* and APOE KO significantly reduce adult neurogenesis (Li *et al*., 2009; Tensaouti et al., 2018; Yang et al., 2011). Therefore, *APOE4’s* influence on adult neurogenesis may be influenced by embryonic neurogenesis.

There are species-specific differences between mouse and human brains, such as increased neuron diversity and ApoE modulation (Maloney et al., 2007; Rakic, 2009). In the human neocortex, outer radial glia (oRG), a population of neural stem cells (NSCs), facilitate a second wave of neurogenesis and give rise to both neurons and glia (Hansen et al., 2010). In the central nervous system, ApoE expression is highest in astrocytes, adult and embryonic NSCs, oRG, and neural progenitors (NPs) (Kim et al., 2009; Yuzwa et al., 2017) Considering that single-cell RNA-sequencing has shown that adult NSC populations resemble embryonic NSCs and oRG, it is possible that the expression of *ApoE4* in oRG, which mice lack, may alter embryonic and adult neurogenesis in similar ways (Baig et al., 2024). To our knowledge, the influence of *APOE4* on embryonic neurogenesis has not been explored in a human-relevant model.

Human induced pluripotent stem cell (iPSC) models have been employed to investigate *APOE4’s* effect on a variety of cell types in the context of AD. *APOE4* increases AD hallmarks Aβ and p-tau in several iPSC-derived neural models (Huang et al., 2022; Lin et al., 2018; Meyer et al., 2019; Wang et al., 2018; Zhao et al., 2020). One study observed that *APOE4* decreased GABAergic neurons, but not glutamatergic neurons, which is consistent with mouse data (Wang *et al*., 2018). In mixed NPs, *APOE4* was associated with accelerated differentiation, maturation, and increased functional neuron maturation (Lin *et al*., 2018; Meyer *et al*., 2019). However, *APOE4*’s influence on embryonic neuron and glial development has not been explored.

We hypothesized that *APOE4* alters neurogenesis and gliogenesis in a neural subtype-specific manner, affecting the maturation and composition of neurons and glia during development. We then reasoned that developmental changes may impact neuronal excitability and network formation. To test our hypothesis, we used human iPSCs to generate regionalized neural organoids patterned towards the cortex (COs) and ganglionic eminence (GEOs) (Birey et al., 2017). We used immunohistochemistry (IHC) at time points aligning with human neurogenic and gliogenic stages. To assess *APOE4*’s functional influence on network formation, we performed three-dimensional (3D) multi-electrode array (MEA) recordings in fused COs and GEOs (assembloids) at later stages.

We discovered that *APOE4* decreases SATB2+ and BRN2+ upper layer neurons and increases astrocytes and oRG during later gliogenic stages, which is associated with increased cell death in COs. In GEOs, *APOE4* accelerates neural differentiation of GABAergic NPs during neurogenic stages, increasing mature neurons at later stages. Lastly, *APOE4* dysregulates GABA-related gene expression in COs, which is associated with altered neuronal network formation and enhanced synchronicity in assembloids. In summary, *APOE4* differentially influences neural subtypes, altering GABA action and network patterns in a manner reminiscent of hyperexcitability. This data supports our hypothesis that *APOE4* influences cortical neural development and has functional repercussions on network formation.

## Results

### *APOE4* promotes gliogenesis while decreasing excitatory neuron subpopulations

To study whether *APOE4* differentially affects the development of neural subtypes, we generated COs and GEOs using a modified protocol adapted from the Pasca lab (Birey *et al*., 2017). Neural differentiation was induced through dual SMAD inhibition to generate COs and, with the addition of ventralizing factors, GEOs from iPSCs (Figure 1A). Given the heterogeneity imposed by genetic background, we used 1 female and 1 male isogenic pair of human iPSCs in addition to 1 control *APOE3/3* (3/3) and 1 AD patient *APOE4/4* (4/4) female lines, totaling 6 iPSC lines with three lines per genotype (Table 1). All iPSC lines expressed pluripotency markers (Figure S1A), had a normal karyotype (Figure S1B), and were confirmed for *APOE* genotype (Figure S1C). Isogenic lines did not harbor any off-target mutations (Nimsanor et al., 2016; Peitz et al., 2018). All iPSC lines were confirmed to be mycoplasma-negative before organoid generation and checked routinely (Figure S1D).

**FIGURE 1.**
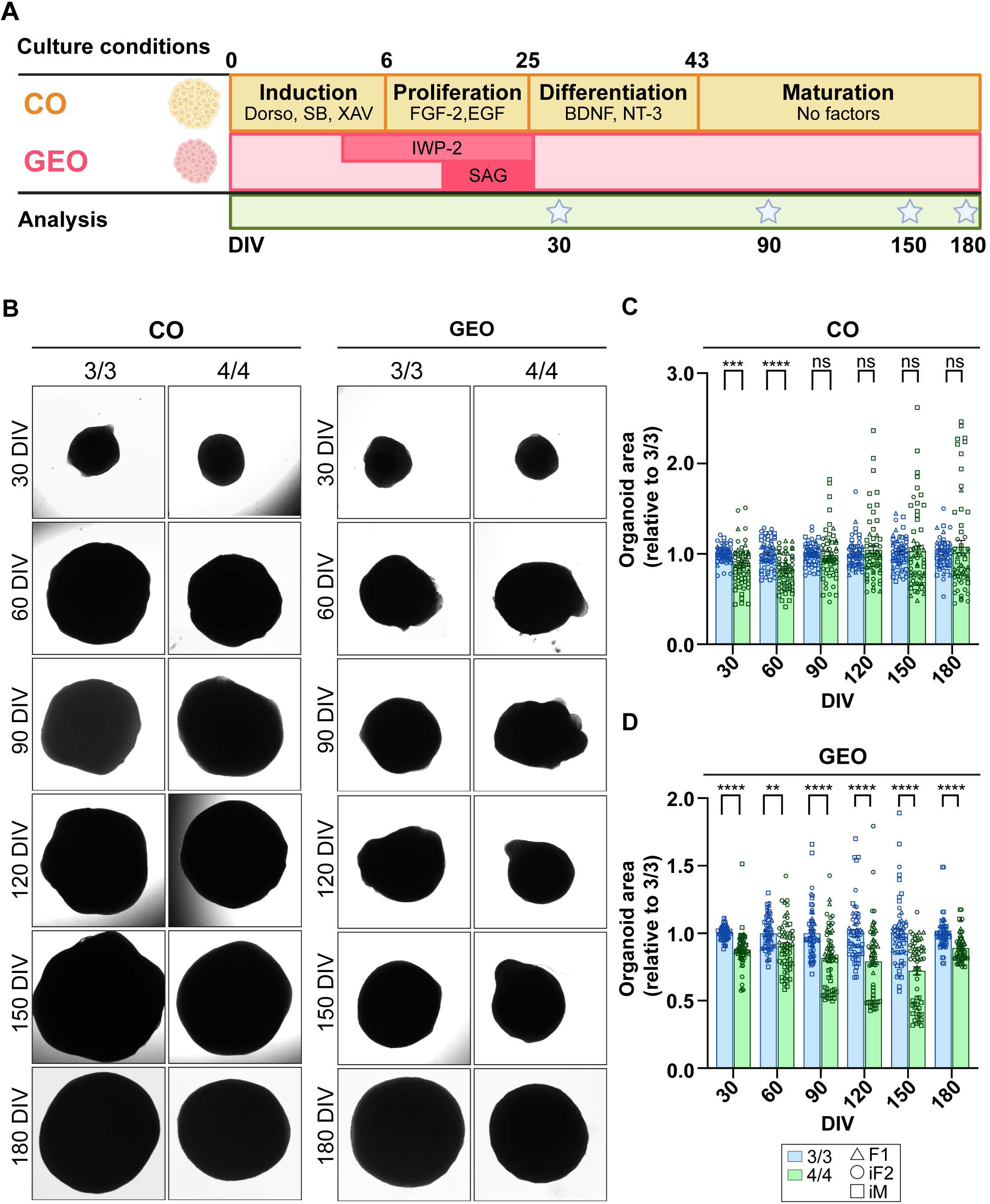
Generation of COs and GEOs from iPSCs and organoid size analysis. **A)** Experimental design to generate COs and GEOs. Organoids were imaged every 30 days from 30 to 180 DIV. Organoids were harvested, and media collected at select time points. **B)** Brightfield representative images of COs and GEOs at all time points. Image area analysis of **C)** COs and **D)** GEOs data with relative size (y-axis) at each time point (x-axis). N= 55-60 organoids from 3 *APOE3/3* and 3 *APOE4/4* iPSC lines (2 independent replicates per iPSC line pair). Data is represented as mean ± SEM relative to *3/3*. Unpaired t-tests with Welch’s correction were utilized to determine significance. ** p < .001, *** p < 0.0005, **** p < 0.0001, ns: not significant. Scale bar = 1 mm.

**Table 1:** human iPSC lines used to generate COs and GEOs.

Manuscript Tables
| Table 1: human iPSC lines used to generate COs and GEOs |  |  |  |  |  |  |  |  |
| --- | --- | --- | --- | --- | --- | --- | --- | --- |
| Paired | Name | Sex | APOE | Age | Clinical Dx |  | Source | Identifier/Source ID |
| F1 | F1 3/3 | F | 3/3 | 75 | Control | Parent | UTSA SCC | 10201-5/ Corriell (AG09173) |
|  | F1 4/4 AD | F | 4/4 | 79 | AD | Parent | CIRM | CW50129FF1 |
| iF2 | F2 3/3 | F | 3/3 | 77 | Control | Parent | EBiSC | BIONi037-A |
|  | F2 i4/4 | F | i4/4 | - | - | Isogenic | EBiSC | BIONi037-A-4 |
| iM | M 4/4 AD | M | 4/4 | 80 | AD | Parent | EBiSC | UKBi011-A |
|  | M i3/3 | M | i3/3 | - | - | Isogenic | EBiSC | UKBi011-A-3 |

To determine whether *APOE4* altered growth during organoid development, COs and GEOs were imaged every 30 days until 180 days *in vitro* (DIV) using brightfield microscopy (Figure 1B). In COs, we observed a size reduction only at 30 and 60 DIV (Figure 1C) while *APOE4* reduced GEO size at all time points analyzed (Figure 1D). This suggests that *APOE4* mediates region-specific alterations in growth, possibly related to cellular composition affecting proliferation, differentiation, and/or survival.

Given the effect of *APOE4* on size, we used IHC to probe COs for differentiation, proliferation, and cell death at the earliest time point. At 30 DIV, COs expressed neural progenitor marker PAX6 and contained ventricular-like regions (Figure S2A) (Nieto-Estévez et al., 2022; Paşca et al., 2015). Ki67, which labels proliferative cells, was expressed in proliferative NSCs and NPs localized within ventricular-like regions (Figure S2A) (Lim et al., 2018). Progenitors then differentiated into immature neurons expressing beta-tubulin-III (TUJ1) (Figure S2A). Cleaved caspase-3 antibody AC3 was used to detect cell death. Using these markers, we saw no difference in proliferation or differentiation in COs (Figure S2B-D). Although cell death levels were generally low, a decrease in cell death was observed in *APOE4* COs at 30 DIV (Figure S2E), possibly due to their smaller size at early time points.

To determine the later effects of *APOE4* on neuronal development in COs, we examined layer II-IV marker SATB2 at 180 DIV, which we found to be decreased in *APOE4* COs (Figure 2A and 2B). A decrease in BRN2, a marker of cortical neuron layers II/III, was observed only in AD *APOE4* paired lines (F1 and iM), suggesting a potential influence of AD genetic background (Figure 2A and 2C). Increased cell death was observed at 180 DIV, with a trending, but nonsignificant, increase observed at 150 DIV (Figure 2A, 2D, S3A, and S3E). At this time point, cell death was not size-dependent, as no significant size difference was observed after 60 DIV (Figure S3A, S3C-E). This data indicates that cell death may contribute to the loss of upper-layer cortical neurons in *APOE4* COs at later developmental stages.

**FIGURE 2.**
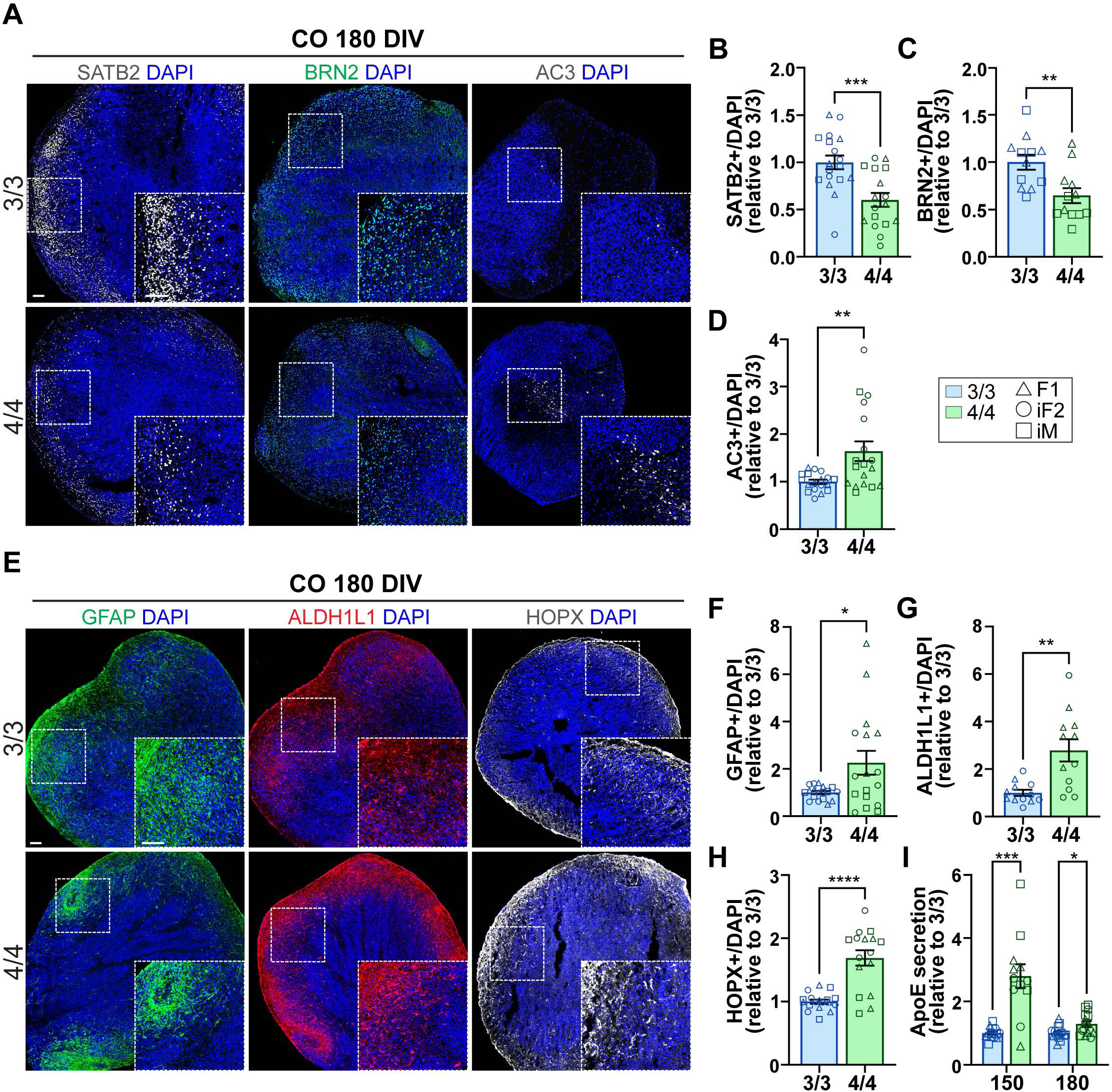
*APOE4* compromises neurogenesis and promotes gliogenesis associated with increased cell death in COs. COs were subjected to IHC for cell type-specific markers and media was subjected to ELISA to measure the levels of secreted ApoE at 180 DIV. **A)** IHC representative images from *APOE3/3* and *APOE4/4* COs for SATB2, BRN2, and AC3. The graphs show quantification of the percentage of **B)** SATB2 **C)** BRN2 and **D)** AC3 over DAPI relative to *3/3*. N= 16-18 organoids from 3 *APOE3/3* iPSC and 3 *APOE4/4* iPSC lines. For BRN2, N=12 organoids from iPSC pairs where *APOE4/4* iPSC is derived from AD patients (F1, iM)**. E)** IHC representative images from *APOE3/3* and *APOE4/4* COs for GFAP, ALDH1L1, and HOPX. Graph quantification of the percentage of **F)** GFAP **G)** ALDH1L1 **H)** HOPX over DAPI relative to *3/3*. **I)** Secreted ApoE levels in COs at 150 and 180 DIV. 1-3 samples of media (each sample from 3-4 organoids) were measured via ELISA. Data is represented as mean ± SEM relative to *3/3*. Unpaired t-tests with Welch’s correction was utilized to determine significance. * p < 0.05, ** p < .01, *** p < .001, **** p < .0001. Scale bar = 100 µm.

An *APOE4-*mediated loss of upper-layer cortical neurons could be attributed to a shift from neurogenesis to gliogenesis. During embryonic development, astrocytes and oRG arise beginning at 3.5 months (aligning with organoid timing) (Amiri et al., 2018; Gordon et al., 2021; Hansen *et al*., 2010). To test whether *APOE4* favors glial differentiation, we measured astrocyte and other glial-specific markers at gliogenic time points (150 and 180 DIV). At 180 DIV, *APOE4* COs showed a significant increase in GFAP, which is primarily expressed in astrocytes at this age (Figure 2E and 2F). Astrocyte markers ALDH1L1 and SOX9 were also increased in *APOE4* COs at 180 DIV (Figure 2E, 2G, S3B and S3F). Additionally, HOPX+ oRG were significantly increased, indicating an overall increase in glia with *APOE4* at gliogenic time points (Figure 2E and 2H). This data suggests that *APOE4* promotes gliogenesis, potentially at the expense of neurogenesis.

Astrocytes and NSCs both express high levels of ApoE while neurons express ApoE in response to pathological conditions, albeit to a lesser extent (Kim *et al*., 2009). To determine the relationship between ApoE secretion and observed phenotypes, we measured secreted ApoE in media by enzyme-linked immunosorbent assay (ELISA) at all time points. At 30-90 DIV, ApoE secretion was below detectable limits likely due to lack of glia and NSCs (data not shown). ApoE was detectable at 120 DIV and all subsequent time points, which aligns with gliogenesis. At 150 and 180 DIV, there was a significant increase in ApoE secretion in *APOE4* COs, further suggesting an increase in astrocytes and oRG, which express the highest amounts of ApoE (Figure 2I).

In the AD brain, altered cleavage of amyloid precursor protein favors cleavage to aggregate-prone Aβ42 over more soluble Aβ40 peptide, resulting in Aβ aggregation. Human iPSC studies in cerebral organoids and cortical neurons observe increased Aβ42/40 ratio and p-tau (Lin *et al*., 2018; Roher et al., 1993; Wang *et al*., 2018). To explore whether COs express AD-related pathologies, we probed total Aβ by IHC, Aβ secretion (Aβ42 and Aβ40) by ELISA, and p-tau by IHC at 180 DIV. Aβ IHC revealed a trending increase in Aβ in *APOE4* COs overall, but we also noted line differences, with significant or trending increases in female *APOE4* COs. (Figure S3G-K). These sex differences also translated to the secreted Aβ42 and Aβ40 and Aβ42/40 ratio. In *APOE4* COs from female lines, Aβ40 secretion was trending or significantly decreased, but Aβ42/40 ratio trended towards an increase. In contrast, the male isogenic line had a highly significant increase in both Aβ42 and Aβ40, but a decrease in Aβ42/40 ratio (Figure S3L-N). No differences were observed between organoid batches (2 replicates per iPSC line). Although COs expressed non-phosphorylated tau robustly, we did not detect any AD-related p-tau at 180 DIV despite attempting several antibodies (data not shown).

Taken together, this data indicates that *APOE4* decreases upper-layer cortical neurons, increases cell death, and increases astrocytes and oRG at developmental time points aligning with the onset of gliogenesis. This suggests that *APOE4* promotes gliogenesis, potentially at the expense of neurogenesis.

### *APOE4* accelerates neuronal differentiation in GEOs

We observed a size deficit in GEOs across all time points; therefore, we sought to determine if *APOE4* affected NP differentiation, maturation, and/or cell death at early and late time points. At 30 DIV, GEOs expressed GE NP marker NKX2.1, which gives rise to GABAergic interneurons. We observed a significant increase in Ki67, NKX2.1, and TUJ1 in *APOE4* GEOs (Figure 3A-D). No difference in cell death was observed (Figure S4A and S4C). These findings suggest that *APOE4* increases GE NP proliferation and differentiation, in line with reports in mixed NPs. Based on these findings, we hypothesized that *APOE4* may accelerate differentiation and/or maturation in GEOs at later time points. At intermediate timepoint 90 DIV, NKX2.1 continued to be elevated in *APOE4* GEOs, suggesting *APOE4* influences GABAergic NP fate decisions (Figure 3E and 3F). At 180 DIV, a trending increase in GABA was observed (Figure 3E and 3G). Calretinin (CR), a calcium-binding protein enriched in a GABAergic neuron subtype, was significantly increased in *APOE4* GEOs at 180 DIV, suggesting that *APOE4* increases neuronal maturation (Figure 3E and 3H). A trending decrease in Ki67 was also observed at 180 DIV, suggesting that *APOE4* neural subtypes were exiting the cell cycle, further supporting increased maturation (Figure S4B and S4F). These data suggest that *APOE4* may accelerate neuronal maturation in GEOs.

**FIGURE 3.**
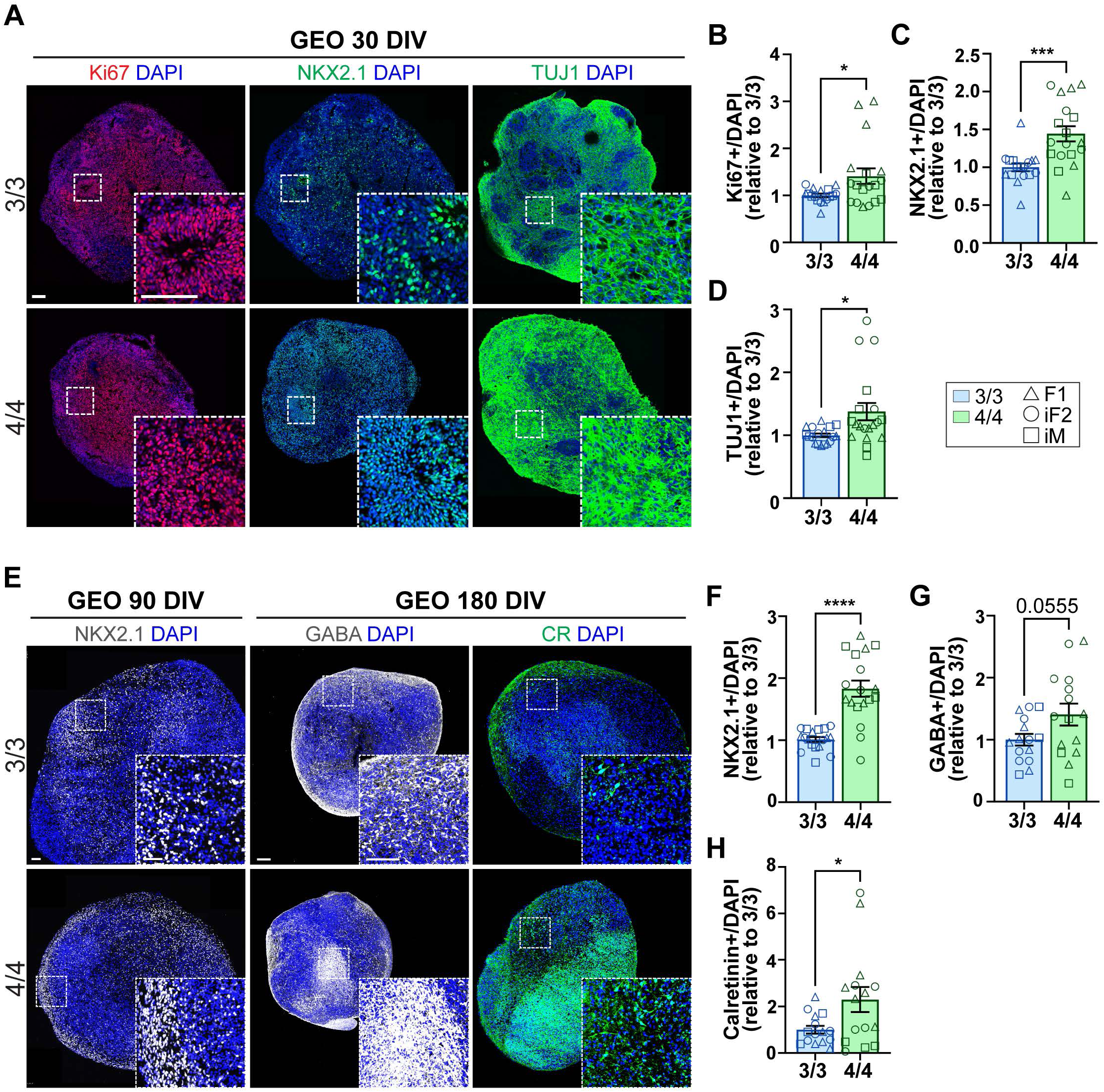
*APOE4* accelerates early neuronal differentiation in GEOs. GEOs were subjected to IHC for cell type-specific markers at 30, 90, and 180 DIV. **A)** IHC representative images from *APOE3/3* and *APOE4/4* GEOs at 30 DIV for Ki67, NKX2.1, and TUJ1. The graphs show quantification of the percentage of **B)** Ki67 **C)** NKX2.1 and **D)** TUJ1 over DAPI relative to *3/3*. **E)** IHC representative images from *APOE3/3* and *APOE4/4* GEOs for NKX2.1 at 90 DIV, GABA, and CR at 180 DIV. The graphs show quantification of the percentage of **F)** NKX2.1 at 90 DIV; and **G)** GABA and **H)** CR over DAPI relative to *3/3* at 180 DIV. N= 16-18 organoids from 3 *APOE3/3* iPSC and 3 *APOE4/4* iPSC lines. Data is represented as mean ± SEM relative to *3/3*. Unpaired t-tests with Welch’s correction was utilized to determine significance. * p < 0.05, ** p < .01, *** p < .001, **** p < .0001. Scale bar = 100 µm.

Of note, there was no difference in ApoE secretion in GEOs, and ApoE secretion was highly dependent on the patient line or isogenic status (Figure S4G), consistent with reports that ApoE secretion and expression correlate with genetic background rather than *APOE* genotype (Tcw et al., 2022). We also note no difference in Aβ42, Aβ40, or Aβ42/40 ratio, and no genotype- or line-specific differences were observed (Figure S4H-J). Similar to COs, there was no expression of p-tau in GEOs despite robust tau expression (Data not shown). Taken together, our data suggests that *APOE4* specifically affects GABAergic NPs by accelerating differentiation and maturation at early time points resulting in increased GABAergic neurons at later time points in GEOs.

### *APOE4* alters neural network activity patterns in fused assembloids

Hyperexcitability is commonly observed in *APOE4* mice and is associated with a loss or impairment of GABAergic neurons (Area-Gomez *et al*., 2020; Gillespie *et al*., 2016; Jang *et al*., 2023; Knoferle *et al*., 2014; Li *et al*., 2009; Nuriel et al., 2017; Palop et al., 2007; Tong et al., 2014; Tong *et al*., 2016). During development, GABA receptors induce excitation due to a higher intracellular chloride ([Cl^-^]_i_) concentration (Ben-Ari, 2002). Within the developing neuronal network, migration, maturation, and synapse formation are modulated by chloride extrusion mechanisms; NKCC1, a Na^+-^K^+^-2Cl^-^ cotransporter is upregulated at early developmental stages, responsible for higher [Cl^-^]_i_, and KCC2, a K^+^- 2Cl^-^ cotransporter is upregulated at later developmental stages, responsible for lower [Cl^-^]_i_ (Ben-Ari, 2002). At 180 DIV, *APOE4* COs, but not GEOs, demonstrated a significant reduction in *GABBR1*, encoding for GABA_B_ receptor subunit 1, (Figure S5A and S5B). We also observed a small yet significant reduction of *SLC12A2* and *SLC12A5*, encoding NKCC1 and KCC2, respectively, in *APOE4* COs (Figure S5C). In *APOE4* GEOs, a trending reduction in *SLC12A2* (NKCC1) and *SLC12A5* (KCC2) was observed (Figure S5D). These findings suggest that *APOE4* may induce a dysregulation of neuronal responsiveness to GABA.

To investigate if *APOE4*-mediated dysregulation of GABA-related genes translates to alterations in neural network formation and function, we utilized a 3D-MEA with 60 electrodes to monitor spontaneous activity. During development, interneurons generated from GE NPs migrate into the cortex where they integrate and form mature cortical neuron circuits (Letinic et al., 2002; Lim *et al*., 2018). To better recapitulate the developing neural network, we fused COs and GEOs (assembloids) at 60 DIV and performed MEA recordings at 200 DIV to assess APOE4’s influence on neuron function (Figure 4A). To confirm GEO interneuron migration into COs, GEO NPs were labeled with a DLX1/2b-GFP expressing lentivirus at 50 DIV, and migration was confirmed at 90-120 DIV (Figure 4B). To explore the functional influence of *APOE4* on GABA action, we applied GABA and picrotoxin (PTX), a GABA_A_ receptor blocker (Figure 4C). Baseline MEA recordings revealed that *APOE4* assembloids exhibited more spontaneous spikes (not significant) with a trending increase in the number of active electrodes (p = 0.07) (Figure 4D and 4E). Network burst analysis revealed 3 out of 4 *APOE4* assembloids had spontaneous network-wide bursts, while only one *APOE3* assembloid did, unless exposed to PTX (Figure 4F). GABA application in *APOE3* assembloids showed minimal impact on spike count or network-wide burst generation, but partially reduced spike count in *APOE4* assembloids (Figure 4D-F). Conversely, PTX enhanced spiking and network-burst activity in all assembloids, with a significantly larger effect in *APOE4*, although this effect is likely due to increased activity overall (Figure 4D and 4F). We additionally assessed the coupling strength between electrodes to gauge the level of synchronous activity within assembloids. Cross-correlogram analysis revealed significantly stronger synchronicity within *APOE4* assembloids compared to *APOE3* in all conditions (Figure 4G). Taken together, our findings indicate that *APOE4*-induced dysregulation of GABA-related gene expression primarily in COs is associated with disrupted network formation resulting in heightened excitability and enhanced synchronicity.

**Figure 4.**
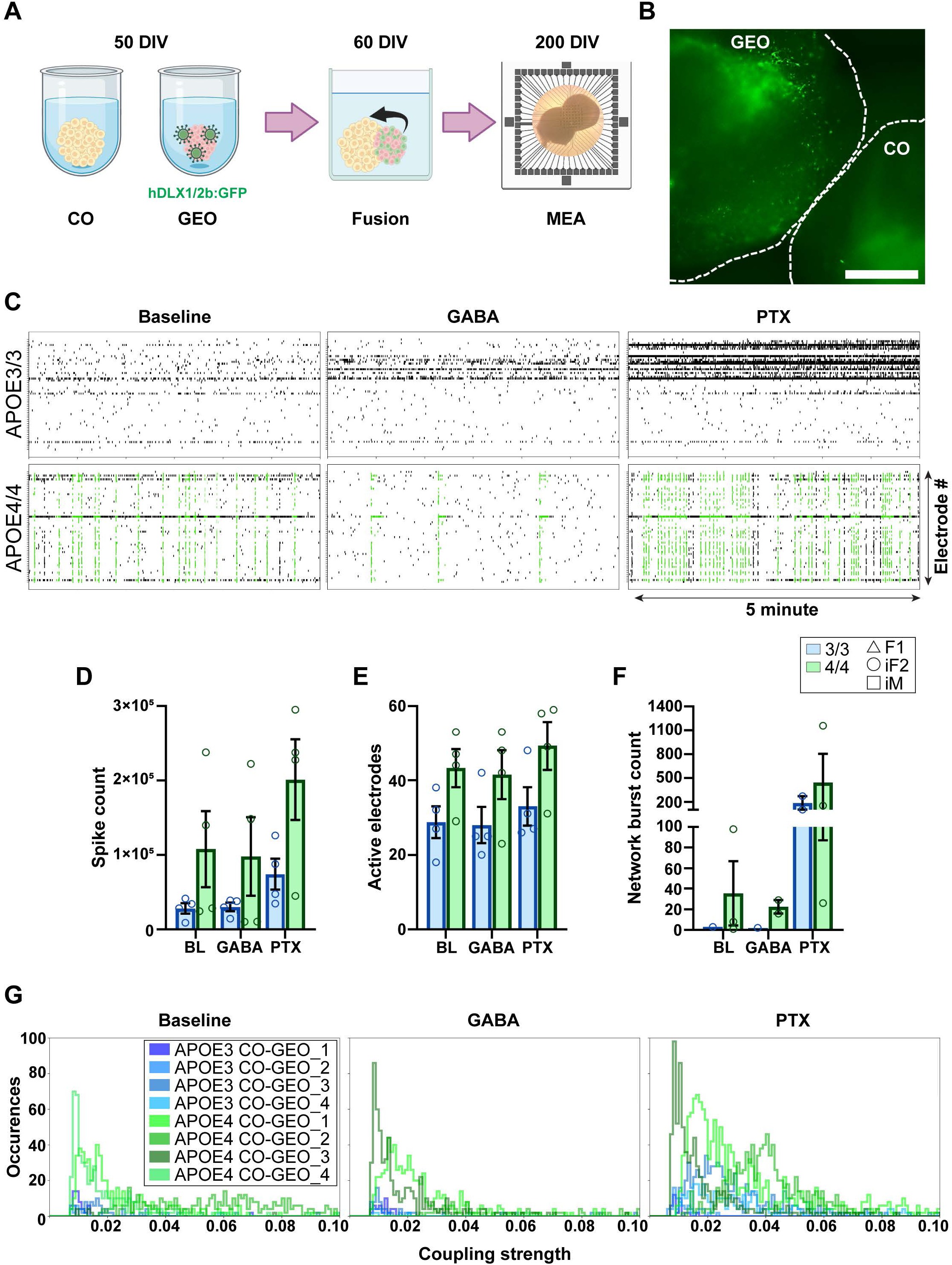
*APOE4* alters neural network activity patterns in assembloids. **A)** Schematic representation of the MEA experiment. At 50 DIV, interneurons were labeled with hDLX1/2:GFP lentivirus in GEOs and fused at 60 DIV. 3D-MEA was performed for electrophysiological assessments of the network at 200 DIV. **B)** At 90 DIV, GFP-labeled interneurons migrate into COs. **C)** Raster plots depict the activity of 60 electrodes (vertical axis) in a representative 5 minutes (horizontal axis) from a 30-minute recording under Baseline, GABA, and PTX administration. The raster plots show increased synchronous activity in *APOE4* assembloids compared to *APOE3*, with green highlighting synchronicity. Bar plots show **D)** the number of spikes **E)** active electrodes and **F)** network bursts during each treatment. **G)** The graphs show the cross-correlation analysis of coupling strength of electrode pairs (horizontal axis) and frequency of coupling strength (vertical axis). N= 4 assembloids from 1 *APOE3/3* iPSC and *APOE4/4* iPSC lines. To determine significance, a 2-way ANOVA with Sidak’s was used for bar plots D-F. Coupling strength refers to the normalized magnitude of cross-correlogram peaks, as detailed in supplemental information. * p < .05, ** p<.01

In summary, *APOE4* differentially affected excitatory and inhibitory cortical neurodevelopment. In COs, *APOE4* resulted in decreased cortical neurons, associated with cell death, and increased glial cell types. In GEOs, we observed increased proliferation and differentiation of GABAergic NPs, resulting in more mature GABAergic neurons. Dysregulation of genes related to GABA action in COs was associated with altered network formation reminiscent of hyperexcitability in *APOE4* assembloids (Figure 5).

**Figure 5.**
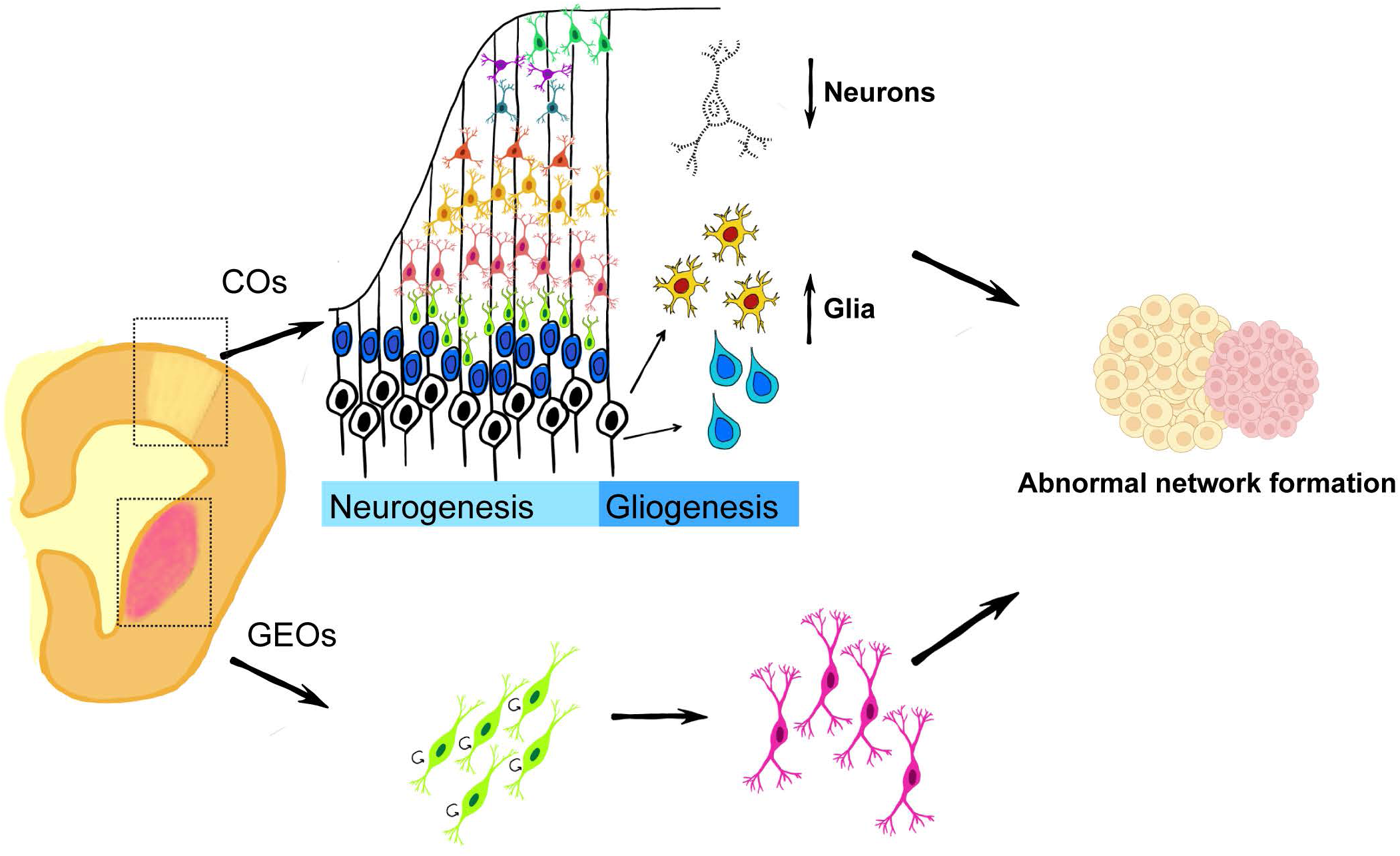
Graphical summary.

## Discussion

*APOE4* induces macroscopic changes in brain structure and cognition in carriers as early as infancy, suggesting that *APOE4* may influence neurodevelopment. While one study observed increased GABAergic neuron vulnerability, most studies focused on mixed NP and neuron populations, and only in the context of AD. The pleiotropic importance of ApoE during neurogenesis and in distinct neural subtypes underscores the need to study *APOE4* in developmental contexts. In the present study, we used regionalized neural organoids from iPSCs to investigate *APOE4’s* influence on excitatory and inhibitory neurons and glia during neurogenic (30-90 DIV) and gliogenic (150+ DIV) stages and explored the functional consequences of *APOE4* on the neural network in assembloids after network establishment (200+ DIV).

We observed size reductions in *APOE4* COs at 30-60 DIV and in GEOs at all time points. In *APOE4* COs, we observed a reduction in BRN2+ neurons in AD *APOE4* iPSC line pairs (F1 and iM) and an overall reduction in SATB2+ neurons, associated with cell death, alongside increased astrocytes and oRG at gliogenic developmental stages. In *APOE4* GEOs, we observed increased NP proliferation and differentiation at 30 DIV resulting in earlier neuronal maturation at 180 DIV. We found dysregulation of gene expression related to GABA action in COs, functionally associated with heightened excitability and synchronicity in assembloids.

### APOE4 decreases cortical layer neurons and increases glia at gliogenic time points

The loss of neurons and increase in glia in COs at gliogenic stages supports the idea that gliogenesis occurs earlier or is favored over neurogenesis with *APOE4*. In mice, *APOE4* decreases adult neurogenesis and increases gliogenesis, suggesting a similar mechanism in embryonic neurogenesis (Li *et al*., 2009). In our human iPSC model, APOE4 dramatically increases oRG (not expressed in mice) and astrocytes, indicating a human-specific phenotype. This is the first study to demonstrate that *APOE4* developmentally alters neurogenic/gliogenic switch, potentially corresponding to macroscopic structural changes in infant APOE4 carriers. Future studies should explore *APOE4’s* influence on oRG function both developmentally and in the context of AD and adult neurogenesis.

### APOE4 accelerates differentiation and maturation in GEOs

At 30 DIV, *APOE4* GEOs demonstrate increased NP proliferation and differentiation via IHC, suggesting a specific influence of *APOE4* on GABAergic NPs in GEOs compared to cortical NPs in COs. In mixed iPSC-derived NPs, increased expression of neurogenesis-related genes (*ASCL1/MASH1*, *DLX2*, and *MEIS1*) was observed in *APOE4* vs *APOE3* (Meyer *et al*., 2019). These genes are involved with GE NP and GABAergic neuron differentiation (Hansen et al., 2013; Kim et al., 2008; López-Tobón et al., 2019; Schmitz et al., 2022). GABAergic NPs might strongly contribute to differential gene expression observed in this study. At later time points, *APOE4* increased GABAergic neuron maturation, as indicated by increased GABA and CR positivity in GEOs. To our knowledge, this is the first publication to independently study excitatory/inhibitory neural subtypes and demonstrate that GABAergic NPs are differentially affected by *APOE4*.

### APOE4 might induce a loss of function during development

A recent study using single-cell RNA-seq found that APOE KO alters neural fate in cerebral organoids, overlapping with our phenotypes (Zhao et al., 2023). They observed decreases in SATB2+ and BRN2+ neurons, increased astrocytes and radial glia, and increased inhibitory neurons. This supports the idea that *APOE4* might induce a loss of function during development. ApoE has a role in adult and embryonic neurogenesis, and adult APOE KO mice show decreased neurogenesis and increased gliogenesis, suggesting an overlap between adult and embryonic neurogenesis (Yang *et al*., 2011). Our data suggest that *APOE4* influences neurodevelopment by reducing ApoE function. Further exploration of *APOE4* on human neurogenesis and oRG might lead to insights into *APOE4’s* role in AD.

### APOE4 results in abnormal neural activity patterns

*APOE4* altered gene expression linked to GABA action in COs at 180 DIV, and increased active electrodes (trending) and synchronicity in assembloids at 200 DIV, suggesting *APOE4* influences network patterning. The increase in activity and synchronicity in *APOE4* assembloids indicates expanded neuronal connectivity and more differentiated neurons and aligns with an observed increase in GABAergic neuron maturation in 180 DIV in GEOs. Despite partial spike inhibition by GABA in *APOE4* assembloids, the network remained highly synchronized, which can be explained by decreased *GABBR1* and KCC2 transporter gene expression in *APOE4* COs (Bassetti, 2022; Wright et al., 2017). *APOE4* assembloids had an increased response to PTX, suggesting variation in maturation levels possibly due to changes in [Cl^-^]_i_ or GABA receptor function. Our findings align with patch-clamp experiments in iPSC-derived *APOE4* neurons revealing larger post-synaptic depolarizing currents (Huang et al., 2019), and possibly contributing to hyperexcitability and synchronization in our *APOE4* assembloids.

### Regionalized organoids express Aβ pathology but not p-tau

We observed an increased trend in Aβ42/40 ratio in female *APOE4* COs. In a previous study from our lab using the same CO protocol, the L435F PSEN1 familial AD (fAD) mutation increased Aβ42/40 and Aβ43/40 ratios at 180 DIV (Hurley et al., 2023). *APOE4* penetrance is not autosomal dominant like fAD mutations, and differences in Aβ may reflect mutation penetrance and severity. Both healthy and AD female *APOE4* carriers exhibit greater AD-related phenotypes than males (Liu *et al*., 2010). We did not detect any p-tau at any time point, although tau was abundant at 180 DIV (data not shown). The lack of p-tau may result from organoid maturity or excitatory/inhibitory neuron interaction. *APOE4* cerebral organoids demonstrate p-tau and Aβ at 12 weeks, while organoid protocols (similar to COs) observe p-tau and Aβ only at 180 DIV suggesting excitatory/inhibitory neuron interactions might induce p-tau (Lin *et al*., 2018; Zhao *et al*., 2020). In conclusion, *APOE4* alters Aβ production and Aβ42/40 ratio, with sex differences, at early developmental stages. Further investigation in different models is needed to understand whether AD pathology is related to early developmental phenotypes.

### Are GABAergic neurons more vulnerable to AD?

In contrast to a study identifying a specific GABAergic neuron loss with *APOE4* in iPSC-derived neurons, we observed a loss of cortical neurons but not GABAergic neurons (Wang *et al*., 2018). These conflicting observations are likely due to technical or activity-dependent differences. A GABAergic vulnerability was demonstrated with a 2D protocol using serum, unlike our protocol. Additionally, in GEOs, when GABA action transitions from excitatory to inhibitory, synaptic firing of GABA inputs onto GABA neurons would result in network-wide inhibition. Therefore, while our reductionist approach allowed us to resolve cell-type specific effects of *APOE4* on neural development, incorporating other cell types might contribute to GABAergic loss. Our findings in COs and assembloids suggest that GABAergic signaling is dysregulated, resulting in network abnormalities. We provide evidence that *APOE4* might mediate cortical neuron vulnerability without necessarily refuting the impact of *APOE4* on GABA signaling impairment. These findings demonstrate a differential influence of *APOE4* on excitatory and inhibitory neurons in a developmental context.

In summary, we provide evidence that *APOE4* influences the development of neurons and glia and may contribute to alterations in network establishment. We provide data demonstrating that *APOE4* differentially affects distinct neural subtypes, with excitatory neurons demonstrating vulnerability, and inhibitory neurons demonstrating increased maturation. *APOE4* resulted in increased astrocytes and oRG generation, which further establishes the possibility that *APOE4* affects neurogenesis and gliogenesis during development in humans. Changes in *APOE4* organoids alter responses to GABA, associated with increased activity and synchronicity. This study is the first to explore APOE4’s influence on the developing human brain using brain organoids. We hope that this study will provide a foundation for further research into APOE4’s influence on human neurodevelopment and its relevance to AD.

## Experimental procedures

### Resource availability

iPSC line 10201-5 was generated and characterized by the UTSA SCC. It is available on request with executed material transfer agreement. Any information and requests for this line should be directed to

### iPSC generation and characterization, and maintenance

See supplemental information for detailed characterization procedures. A total of 6 human induced pluripotent stem cell (iPSC) lines were sourced for this study (Table 1). All iPSC lines had normal karyotypes (Wicell) and expressed pluripotency markers (Lin28, Nanog, Oct3/4, and SOX2). iPSCs were maintained in mTeSR™ 1 medium (Cat. No. 05851, Stemcell Technologies) on 6-well tissue culture plates (Cat. No. 3506, Corning) coated with growth factor reduced Matrigel (Cat. No. 356230, BD Biosciences). Upon thawing, ROCK inhibitor Y27632 (final concentration 10 µM, Cat. No. S-1049, Selleck Chemicals), was added. Cells were passaged at 70% confluence, and Versene solution (Cat. No. 15040-066, Thermo Fisher Scientific) was used to detach cells for replating at 1:12 or frozen in knockout™ serum replacement (Cat No. 10828028, Gibco) and 10% DMSO.

### Genotyping

APOE genotype was confirmed by Sanger sequencing (Eurofins Genomics LLC). Genomic DNA was extracted from iPSCs using DNeasy® Blood & Tissue Kit (QIAprep #69504) following the manufacturer’s instructions. OneTaq® Hot Start DNA Polymerase (New England BioLab Cat. No. M0481) was used to amplify the product containing the 2 basepairs that differ between APOE alleles using primers (Forward CTGGAGGAACAACTGACCCC, Reverse CTCGAACCAGCTCTTGAGG). PCR master mix was used according to the manufacturer’s instructions with the addition of 7.5% DMSO. PCR conditions were as follows, 94 °C for 4 minutes; 94 °C 30 sec, 65 °C 45 sec, 68 °C 1 minute for 40 cycles; 68 °C for 5 minutes. PCR products (∼550 bp) were visualized on an agarose gel, purified (QIAquick PCR Purification Kit Cat. No. 28106), and sent for Sanger sequencing.

### CO and GEO generation

Organoids were generated using the methods described by Pasca and colleagues adapting slight modifications (Birey *et al*., 2017; Sloan et al., 2017). More detailed culture conditions, reagents, and media formulations can be found in supplemental information. Briefly, on day 0, iPSCs were seeded at 9,000 cells per well of a 96-well round bottom plate in mTeSR containing 20 µM of Y27632. On days 1-5, Dual-SMAD inhibition was achieved with 2.5 µM of dorsomorphin and 10 µM of SB-431542, and 1.2 µM of wnt-inhibitor XAV 939 to enhance forebrain differentiation in TeSR™-E6 medium. Neural medium was used after day 6 for the rest of the experiment. On days 6-24, 20 ng/ml of bFGF and 20 ng/ml of EGF were added to neural medium to promote proliferation. Organoids were transferred to 24 well plates on day 17 and were maintained on an orbital shaker to promote oxygenation. On days 25-42 20 ng/ml of BDNF and 20 ng/ml of NT-3 were added to promote differentiation. For GEO generation, 5 µM of Wnt inhibitor IWP-2 was added on days 4-22, and 100 nM of SMO pathway activator SAG was added on days 12-22. Each cell line pair (*APOE3/3* and *APOE4/4* isogenic pair, or *APOE3/3* control and *APOE4/4* AD patient line) was differentiated into organoids two independent times (experimental replicate), and for IHC or qRT-PCR 3 organoids were collected for each time point for each experimental replicate.

### IHC and sample preparation

**For iPSCs:** iPSCs were plated on glass coverslips in a 24-well plate. When cells reached 60-70% confluence, they were fixed with 4% paraformaldehyde (PFA) for 15 minutes and washed 3 times with PBS. Coverslips were stained using the same protocol as described below for organoids. **For organoids:** At each time point, organoids were harvested and fixed in 4% PFA for 1-2 hours at room temperature or overnight at 4 °C, then incubated at 4 °C in 30% sucrose until they sank (24-48 hours). Organoids are embedded in OCT compound and frozen on dry ice. Organoids were cryosectioned at 14-µm, in serial, on glass slides. For IHC, organoids were incubated in blocking buffer (1X Carbo-Free Blocking solution (Vector labs, Cat. No. SP-5040-125), 0.3% Triton X-100 in TBS) for 1 hour at room temperature (RT), and primary antibodies (See Table S1) overnight at 4 °C in a humidified chamber. After 3 TBS washes, slides were incubated with secondary antibodies (Jackson Immunoresearch, 1:400, or Alexafluor 488, 1:1000, or Alexafluor 647, 1:400) for 2 hours at RT. Slides were washed, 4’,6-diamidino-2-phenylindole (DAPI; Sigma, Cat. No. D9542) was added to the second wash to label nuclei, then coverslipped using polyvinyl Alcohol solution (PVA; Sigma, Cat. No. BP168-122). Four sections per organoid were imaged using a Leica (Spe8-II), or Nikon (A1R HD25) confocal microscopes, and fluorescence intensity was analyzed using Image J software. For analysis, the thresholded area of each marker was divided over thresholded DAPI for nuclear markers, and organoid area for cytoplasmic markers. 4 sections were analyzed per organoid with a total of 3 organoids per replicate (2 replicates per iPSC line pair, 3 iPSC line pairs, N=18 organoids). Data is represented relative to APOE3/3 for all IHC.

### RNA isolation, Reverse Transcriptase (RT) reaction, and quantitative PCR assay

Organoid RNA was extracted using the Qiagen miRNeasy Mini Kit (Cat. No. 217004) according to manufacturer instructions. The concentration and purity of the RNA samples were measured by using Nano-drop (Thermo Fisher Scientific). The extracted RNA (500 ng) was reverse transcribed according to the protocol supplied with SuperScript III First-Strand Synthesis System for RT (Invitrogen, Cat. No. 18080-051). Quantitative real-time PCR (qRT-PCR) was carried out in a QuantStudio5 real-time PCR system using the PowerUp SYBR Green Master Mix methodology following the manufacturer’s instructions (Applied Biosystems, Cat. No. A25742). Reactions were run in triplicate and the expression of each gene was normalized to the geometric mean of GAPDH as a housekeeping gene and analyzed by using the ΔΔCT method. The primer sequences of each gene are listed in Supplemental Table 2.

### Measurement of ApoE, Aβ_40,_ and Aβ_42_ in the medium from organoids

At selected time points, the medium was collected from organoid cultures (3-4 organoids after 3-4 days in media) and stored at -80 °C. Aβ peptides were measured with Human β Amyloid (1-42) ELISA kit (Wako Chemicals, Cat. No 298-624-01) with undiluted media and Wako Human β Amyloid (1-40) ELISA kit (Wako Chemicals, Cat. No 298-64601) with 1:4 diluted media. APOE media was measured with Apolipoprotein E Human ELISA Kit (Thermo Fisher Scientific, Cat. No. EHAPOE) with 1:3 diluted media. Plates were measured with GloMax® Explorer Multimode Microplate reader (GM3500), and data were analyzed using Graphpad Prism 9 software.

### MEA recording and analysis

Detailed protocol, equipment, and analysis used can be found in supplemental information. Organoids were maintained using BrainPhys^TM^ hPSC Neuron kit (05795; STEMCELL Technology) for a week before and during recording sessions. All MEA experiments were performed on 200 DIV organoids using 3D-MEA (60-3DMEA200/12/80iR-Ti, Multichannel system, Harvard Bioscience) chips. Sequential recordings were performed as follows: baseline, 20 µM GABA (56-12-2, Sigma-Aldrich), and 58 µM picrotoxin (124-87-8, Sigma-Aldrich), for 30 minutes each condition, with recording started 10 minutes after drug application. Raw electrical potentials were amplified using an electronic amplifier (ME2100-Mini, Multichannel systems, Harvard Bioscience), sampled at 25 kHz/channel, and digitized at 16-bit resolution. A peak detection algorithm with an adaptive threshold was employed for spontaneous network-wide synchronization of spike times (i.e. network bursts) (Mahmud et al., 2014; Quiroga et al., 2004). The events were visualized as a raster plot and further analyzed by spike train analysis (Mahmud *et al*., 2014; Quiroga *et al*., 2004). Interaction between electrode pairs was derived using cross-correlation analysis of spike times (Knox, 1981). Peak value distribution across experimental conditions was generated to facilitate comparisons of coupling strength. For baseline spike count and active electrodes, a two-tailed unpaired t-test was used to determine significance. A paired t-test was used to compare the effect of chemical application and genotype on active electrodes and spike count.

### Statistical analysis

Outliers were removed from each dataset using Graphpad Prism software (Q=1%). A two-tailed unpaired Student’s t-test was used to compare the mean ± standard error of the mean (SEM) values, with Welch’s correction when the F-test indicated significant differences between the variances of both groups. For data shown as relative to control, all values were normalized to the average of the control within each replicate. With the exception of coupling strength analysis, all analyses were carried out with GraphPad Prism software and the differences were considered statistically significant when P < 0.05. For MEA bar graphs, a paired t-test was used to compare the effect of chemical application and genotype (within each of the organoids) on active electrodes and spike count. Coupling strength analysis was performed using MATLAB (See supplementary information for details).

### Declaration of generative AI-assisted technologies in the writing process

During the editing process, the authors used ChatGPT to reduce word count and redundancy without changing the manuscript structure or content. All minor changes were curated by the authors, and we take full responsibility for the content of the publication.

## Acknowledgments

This work was supported by NIH grants (U01DA054170, R01NS113516, R01NS124855, and R21AG066496) and the Robert J. Kleberg, Jr. and Helen C. Kleberg Foundation and the Semmes Foundation (to J.H.); and NIH 1F31AG082498 (to K.M.-A.). We would like to thank Bess Frost, Hyoung-gon Lee, and Chris Gamblin for their help with reagents, protocols, and advice on the project. Some figures were created with BioRender.com.

## Author contributions

Conceptualization, K. M.-A., J.H. and V.N.-E.; Methodology, K. M.-A., E.D.-G. and J.H.; Software, A.H.; Formal Analysis, K.M.-A., E.D.-G., S.M., and A.H.; Investigation, K.M.-A., E.D.-G., A.A., S.M., A.M.P., and Y.R.; Resources, C.N.; Writing-original draft, K.M.-A.; Writing-review & editing, K.M.-A., E.D.-G, S.M., A.M.P., V.N.-E., and J.H.; Visualization, K.M.-A.; Supervision, K.M.-A., M.G., and J.H.; Funding Acquisition, K.M.-A. and J.H.

