## Supplemental information for "*APOE4* impacts cortical neurodevelopment and alters network formation in human brain organoids"

Figure S1 - iPSC characterization

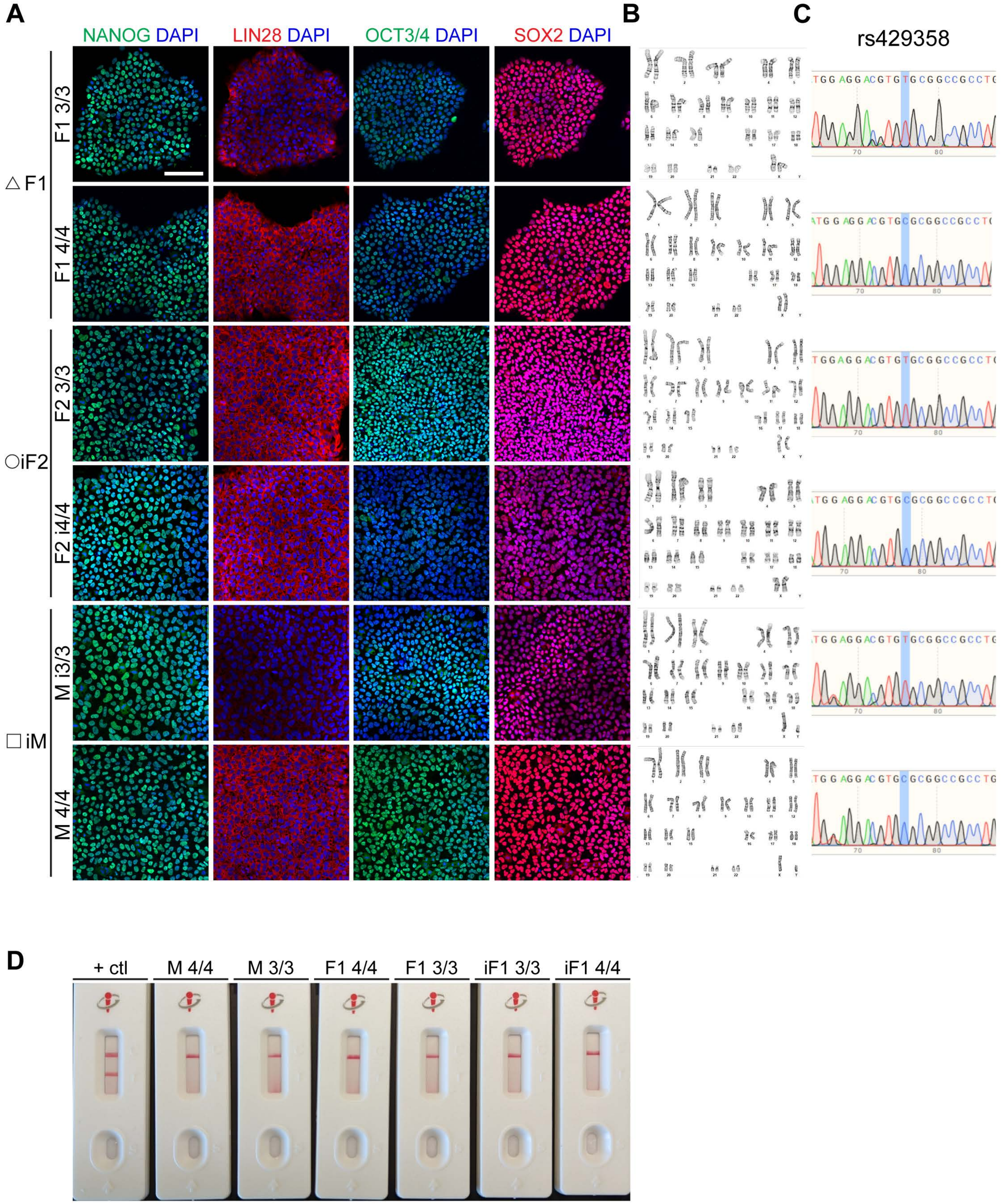

**Figure S1. iPSCs characterization. A)** ICC representative images of each iPSC line used in this study showing the expression of the pluripotency markers Nanog, Lin28, OCT3/4, and SOX2. The nucleus was stained with DAPI. Each line was confirmed to have **B)** a normal karyotype, and **C)** a correct genotype. **D)** representative image of routine mycoplasma testing negativity, which are performed prior to and routinely during organoid culture. Scale bar = 100 μm.

Figure S2 - CO 1

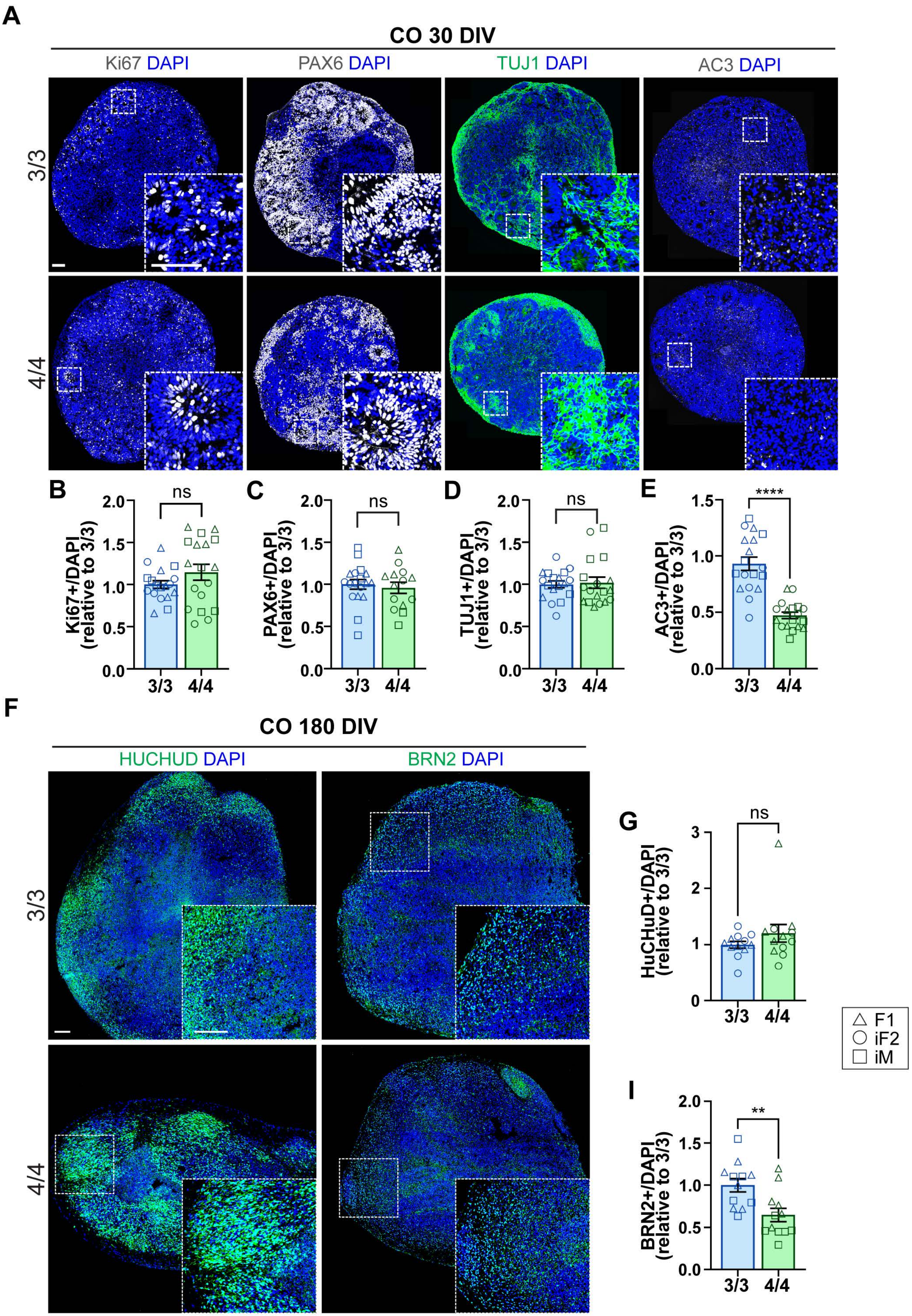

**Figure S2. Characterization of COs at early time points.** COs were subjected to IHC for further characterization of cell type-specific markers and cell death at 30 DIV. **A)** IHC representative images of *APOE3/3* and *APOE4/4* COs immunoassayed for Ki67, PAX6, TUJ1, and AC3. The graphs show quantification of the percentage of **B)** Ki67 **C)** PAX6 **D)** TUJ1 and **E)** AC3 positivity/DAPI normalized to *APOE3/3*. N= 16-18 organoids from 3 *APOE3/3* iPSC and 3 *APOE4/4* iPSC lines. Data were normalized to *APOE3/3* for each replicate (2 replicates per iPSC line). Data is represented as mean ± SEM relative to 3/3. Unpaired t-tests with Welch's correction were utilized to determine significance. * $p < 0.05$, **** $p < .0001$, ns: not significant. Scale bar = 100 μm.

Figure S3 - CO 2

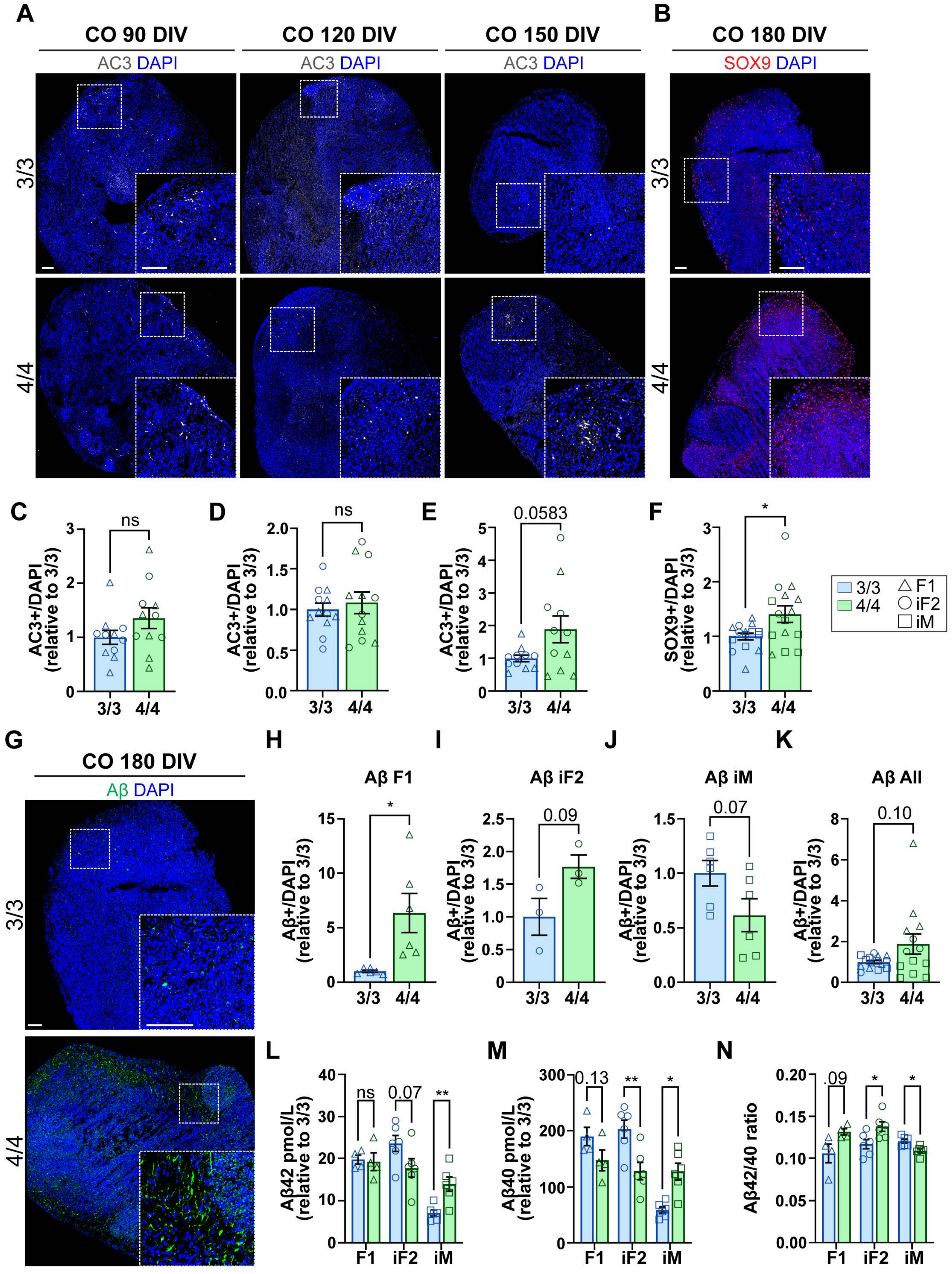

**Figure S3. Characterization of COs at intermediate and late time points.** COs were subjected to IHC for further characterization of cell type-specific markers, cell death, and Aβ pathology at intermediate and late time points. Media was subjected to ELISA to measure the levels of secreted Aβ42 and Aβ40 at 180 DIV. IHC representative images of *APOE3/3* and *APOE4/4* COs at **A)** 90, 120, and 150 DIV immunoassayed for AC3, and **B)** SOX9 at 180 DIV. Under each IHC image is the respective graph quantification of the percentage of AC3 at **C)** 90 **D)** 120 and **E)** 150 DIV represented relative to 3/3 and **F)** SOX9 represented relative to 3/3. **G)** IHC representative images of *APOE3/3* and *APOE4/4* COs at 180 DIV immunoassayed for Aβ (D54D2). The graphs show the quantification of Aβ in **H)** F1 **I)** iF2 **J)** iM and **K)** all together represented relative to 3/3. The graphs show the secreted levels of **L)** Aβ42 and **M)** Aβ40 separated by iPSC line measured by ELISA from media collected from COs at 180 DIV. **P)** The graph shows the Aβ42/Aβ40 ratio separated by line. 90-150 DIV: N= 12 organoids from 2 *APOE3/3* iPSC and 2 *APOE4/4* iPSC lines. 180 DIV: N= 16-18 organoids from 3 *APOE3/3* iPSC and 3 *APOE4/4* iPSC lines. 1-3 samples of media (media from 3-4 organoids per sample) were measured via ELISA. Data is represented as mean ± SEM relative to 3/3. Unpaired t-tests with Welch's correction was utilized to determine significance. * $p < 0.05$ ns: not significant. Scale bar = 100 μm.

Figure S4 - GEO

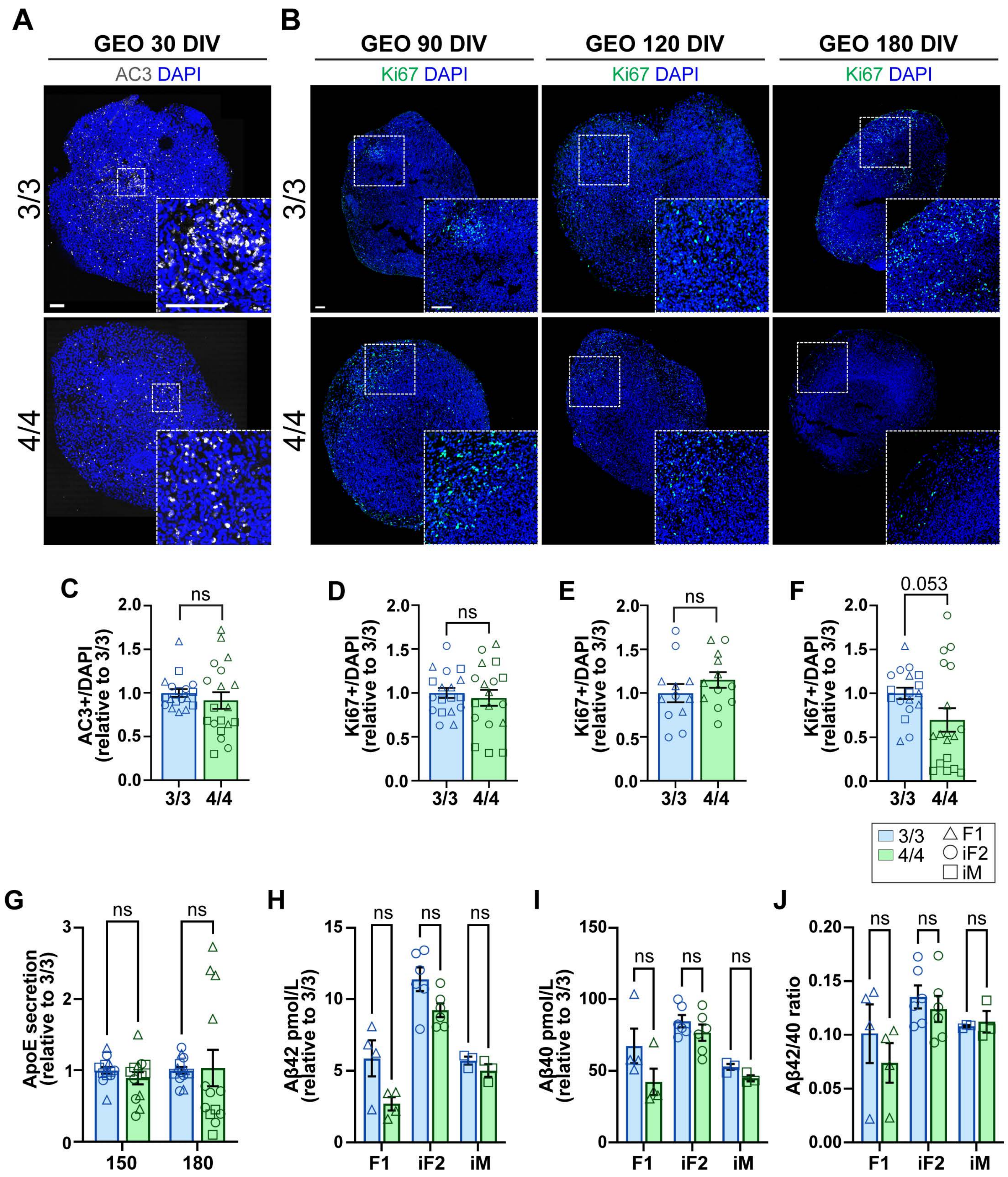

**Figure S4. Characterization of GEOs**. GEOs were subjected to IHC for further characterization of cell death and proliferation at early and intermediate time point, and media was subjected to ELISAs for measuring the levels of secreted ApoE, Aβ42, and Aβ40 at late time points. **A)** IHC representative images of *APOE3/3* and *APOE4/4* COs at 30 DIV immunoassayed for AC3. **B)** IHC representative images of *APOE3/3* and *APOE4/4* COs immunoassayed for Ki67 at 90, 120, and 180 DIV. Under each IHC image is the respective graph quantification of **C)** AC3 at 30 DIV and Ki67 at **D)** 90 **E)** 120 and **F)** 180 DIV over DAPI relative to *3/3*. **G)** The graph shows the levels of secreted ApoE in GEOs at 150 and 180 DIV measured by ELISA represented relative to 3/3. The graphs show the secreted levels of **H)** Aβ42 and **I)** Aβ40 separated by iPSC line measured by ELISA from media collected from GEOs at 180 DIV. **J)** The graph shows the Aβ42/Aβ40 ratio. IHC: N= 16-18 organoids from 3 *APOE3/3* and 3 *APOE4/4* iPSC lines. Data is represented as mean ± SEM relative to 3/3. ELISA: 1-3 samples of media (3-4 organoids per sample) were measured. IHC and ApoE ELISA data were normalized to *APOE3/3* for each replicate (2 replicates per iPSC line). Data is represented as mean ± SEM. Unpaired t-tests with Welch's correction was utilized to determine significance. ns: not significant. Scale bar = 100 μm.

Figure S5 - GABA and assembloids

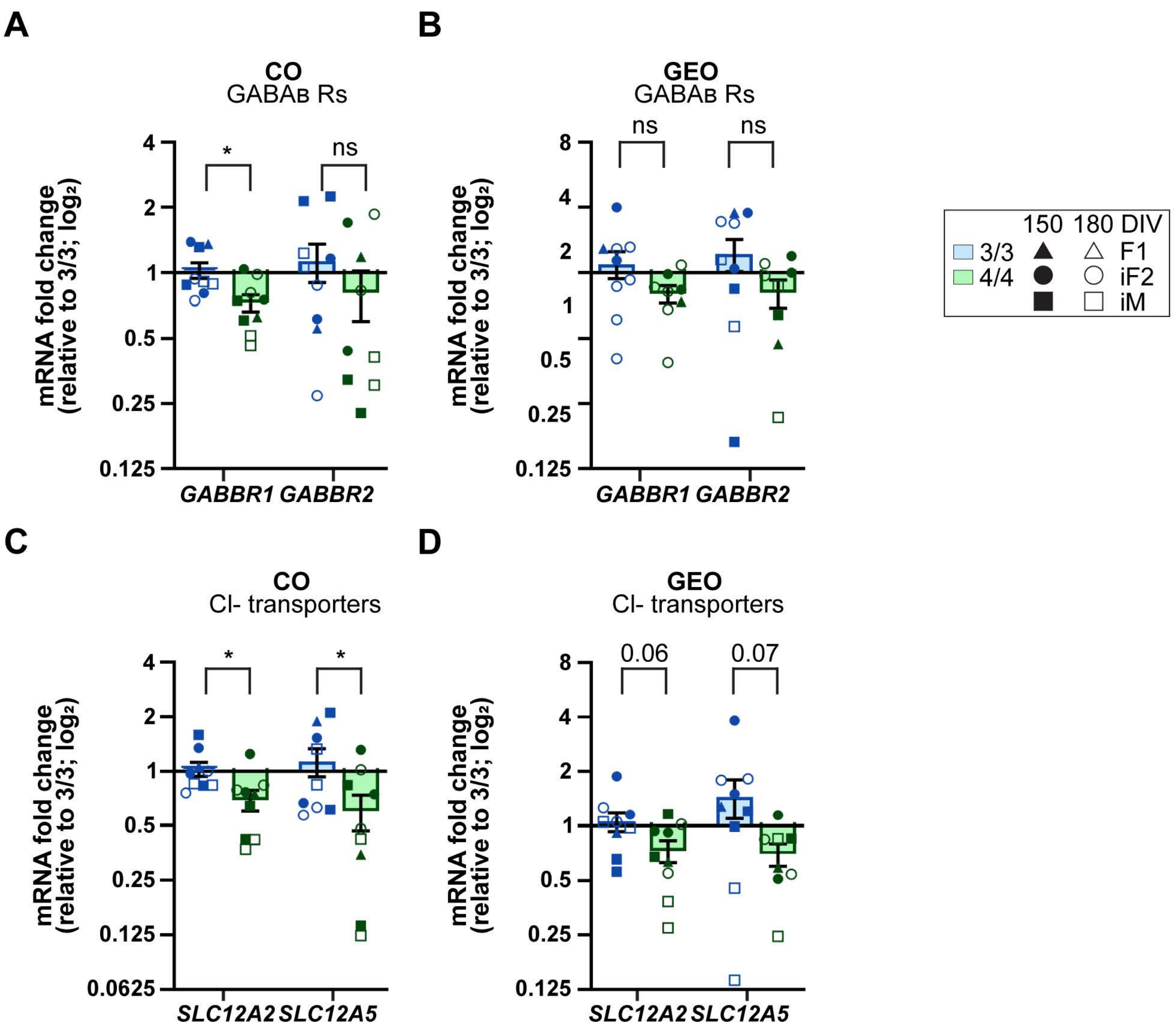

**Figure S5. GABA-related gene expression in COs and GEOs at 180 DIV.** At 150 and 180 DIV, organoids were harvested for gene expression analysis. The expression levels of $GABA_B$ receptors and $Cl^-$ transporters were quantified by qRT-PCR using SYBR green assay. The graphs show the fold change in the expression of $GABA_B$ receptors *(GABBR1* and *GABBR2 genes*) in **A)** *APOE4* COs and **B)** *APOE4* GEOs compared to *APOE3*; and the fold change in the expression of $Cl^-$ transporters *(SLC12A2 and SLC12A5 genes*) in **C)** *APOE4* COs and **D)** *APOE4* GEOs compared to *APOE3*. N= 9 samples from 3 *APOE3/3* and 3 *APOE4/4* iPSC lines (3 organoids per sample; 2 replicates per iPSC line). Data is represented as mean ± SEM. Unpaired t-test was utilized to determine significance. * $p < 0.05$, ns: not significant.

#### Supplemental Tables

| Table S1: List of primary antibodies | | | | |
|---|---|---|---|---|
| **Antibody** | **Host Species** | **Company** | **Cat. Number** | **Dilution** |
| **Anti-AC3** | Rabbit | Cell Signaling | 9661 | 1:400 |
| **Anti-ALDH1L1** | Rabbit | Abcam | Ab177463 | 1:200 |
| **Anti-BRN2** | Mouse | EMD Millipore | MABD51 | 1:50 |
| **Anti-Calretinin** | Mouse | Swant | CR6B | 1:1000 |
| **Anti-D54D2 (Aβ)** | Rabbit | Cell Signaling | 8243 | 1:500 |
| **Anti-GABA** | Rabbit | Sigma | A2052 | 1:1000 |
| **Anti-GFAP** | Chicken | Millipore Sigma | AB5541 | 1:1000 |
| **Anti-HucHuD** | Mouse | Invitrogen | A-21271 | 1:500 |
| **Anti-HOPX** | Rabbit | Sigma | HPA055888 | 1:500 |
| **Anti-Ki67** | Mouse | BD | 550609 | 1:500 |
| **Anti-Ki67** | Rat | Invitrogen | 14-5698-82 | 1:200 |
| **Anti-LIN28** | Rabbit | Cell Signaling | 3978 | 1:1000 |
| **Anti-MAP2ab** | Mouse | Sigma | M1406 | 1:500 |
| **Anti-NANOG** | Mouse | Thermoscientific | MA1-017 | 1:500 |
| **Anti-NKX2.1** | Rabbit | Abcam | ab133737 | 1:500 |
| **Anti-Oct3/4** | Mouse | Santa Cruz | sc-5279 | 1:1000 |
| **Anti-PAX6** | Rabbit | Biolegend | 901301 | 1:300 |
| **Anti-SATB2** | Rabbit | Abcam | AB34735 | 1:500 |
| **Anti-SOX2** | Rabbit | Millipore | AB5603 | 1:1000 |
| **Anti-SOX9** | Goat | R&D Systems | AF3075 | 1:500 |
| **Anti-TUJ1** | Mouse | Sigma | T8660 | 1:400 |

| Table S2: List of primers for RT-qPCR | | |
|---|---|---|
| **Gene** | **Forward** | **Reverse** |
| **GABBR1** | TTCAACTACAACAACCAGACCATTACCG | GCGTCCATGCCATCCGAGAG |
| **GABBR2** | CCCCTGCGAAGGACAGTGGAG | AACAACCGAACAACATGAGAAGTCCC |
| **GAPDH** | GGAAGCTTGTCATCAATGGAAATC | TCAGCAGAGGGGGCAGAGAT |
| **SLC12A2** | AACGCTGTTGCAGTTGCTATGTATGTG | AGATACCTAAAAGAATCACGACTGTAATGGC |
| **SLC12A5** | CTGCAGAACATCTTTGGCGTCATC | CAGCAGGCACAACACCATTCGT |

### Supplemental experimental procedures

##### iPSC characterization continued:

*APOE4/4* female iPSC line (CW50129) was sourced from CIRM. An iPSC line was generated by the UTSA stem cell core from fibroblast samples (Coriell) using Cytotune-iPS Sendai Reprogramming kit (Thermofisher). We also confirmed no integration of reprogramming genes. All iPSCs were expanded to create stocks from 3-4 passages then karyotyped (Wicell), and experiments were performed within 5 passages of karyotype. For pluripotency, iPSC's were grown on coverslips in a 24-well plate and were fixed at 75% confluence for immunocytochemistry (see below). All iPSC lines were negative for mycoplasma throughout the study using either genomic PCR (Cat No. MP0035-1KT, Millipore sigma) or supernatant (Cat No. rep-mys-20, Invitrogen). iPSCs were maintained without antibiotics with no visible signs of microbial presence.

##### Detailed organoid generation protocol

iPSCs were seeded at 275,000 cells per well of a 6-well plate to reach 70% confluence at the appropriate time. 12-24 hours prior to organoid generation, 1% DMSO was added during routine feeding to enhance the differentiation of iPSCs (Chetty et al., 2013). On day 0 of differentiation, iPSCs at 70% confluence were detached and dissociated with Accutase (Sigma-Aldrich) and washed with mTESR. iPSCs were seeded at 9,000 cells per well of an ultra-low attachment 96-well round bottom plate (Corning Cat. No. 7007) in mTeSR containing Y27632 (20 μM), and medium changes were performed daily. On Day 1, media was changed to neural induction medium TeSR™-E6 medium (Cat. No. 05946, Stemcell Technologies). On days 1-5, Dual-SMAD inhibition was performed using SMAD inhibitors dorsomorphin (2.5 μM, Sigma) and SB-431542 (10 μM, Tocris), and wnt-inhibitor XAV 939 (1.2 μM, Tocris) was added to enhance forebrain differentiation. On days 6-24, media was replaced with neural medium (Neurobasal A, B-27 without Vitamin A, Glutamax, penicillin/streptomycin) containing the growth factors, bFGF (20 ng/ml, Peprotech) and EGF (20 ng/ml, Peprotech). On day 15, organoids were transferred to a 24-well plate (Corning Cat. No. 3473) and were maintained on an orbital shaker (80 RPM) to promote oxygenation. On day 25, media was replaced with neural medium containing BDNF (20 ng/ml, Peprotech) and NT-3 (20 ng/ml, Peprotech) until day 42. For ganglionic eminence organoid generation, Wnt inhibitor IWP-2 (5 μM, Selleckchem) was added on days 4-22, and the SMO pathway activator, SAG (100 nM, Selleckchem) on days 12-22.

##### Detailed MEA recording and analysis:

All recordings were carried out inside a humid (i.e., 95% R.H.) incubator at 37℃ and 5% $CO_2$. Organoids were maintained using BrainPhys$^{TM}$ hPSC Neuron kit (05795; STEMCELL Technology) for a week prior to recording session. For each session, individual organoids were placed on 3D – MEA (60-3DMEA200/12/80iR-Ti, Multichannel system, Harvard Bioscience). To ensure optimal attachment to the MEA surface, organoids were immersed in 200 μL of BrainPhys$^{TM}$. Sequential recordings were performed as follows: baseline, 20 μM GABA (56-12-2, Sigma-Aldrich), and 58 μM picrotoxin (124-87-8, Sigma-Aldrich), for 30 minutes each condition, with recording started 10 minutes after drug application. Fluorinated Teflon thin film (MEA-MEM-set5, Multichannel systems, Harvard Bioscience) was used to seal the chip, to block evaporation and contamination. Raw electrical potentials were amplified using electronic amplifier (ME2100-Mini, Multichannel systems, Harvard Bioscience), sampled at 25 kHz/channel, and digitized at 16-bit resolution. Data were stored on disk via software (Experimenter, Multichannel systems, Harvard Bioscience). To detect the time of occurrence of putative action potentials (i.e. spike times), a peak detection algorithm with adaptive threshold was employed, for spontaneous network-wide synchronization of spike times (i.e. network bursts) could be detected and quantified (Mahmud et al., 2014; Quiroga et al., 2004). The events were visualized as a raster plot and further analyzed by spike train analysis (Mahmud *et al.*, 2014; Quiroga *et al.*, 2004). Interaction between electrode pairs was derived using cross-correlation analysis of spike times (Knox, 1981), limited to inter-spike delays less than 500 ms and quantified by 3 ms bins. Each cross correlogram was divided by the square root of spike number in a spike train to account for firing rate modulation. The peak value from the cross-correlogram represented connectivity strength of the electrode pairs. Peak value distribution across experimental conditions were generated to facilitate a comparison of coupling strength. To assess the significance of these peaks (coupling strength), their corresponding inter-spike intervals (ISI) were randomly shuffled to achieve surrogate spike times with identical distribution of ISI. The peaks were considered significant if they showed values larger than the mean plus 3 standard deviations of the cross-correlogram of the surrogate.
